# Metatranscriptomic surveillance reveals overlooked liver-associated parasite diversity and broad host ranges in small mammals from southern China

**DOI:** 10.64898/2026.09.12.751130

**Authors:** Min-Wu Peng, De Wu, Qing-Quan Cen, Qiang Mao, Yu-Qi Liao, Geng-Yang Xin, Li-Jun Chen, Jia-Hui Liu, Wen-Cheng Lu, Jing-Diao Chen, Mang Shi

**Affiliations:** Institute of Parasitic Diseases, Center for Disease Control and Prevention of Guangdong Province, Guangzhou, China; Zhanjiang Institute of Plague Control and Prevention, Zhanjiang, China; State Key Laboratory for Biocontrol, School of Medicine, Shenzhen Campus of Sun Yat-sen University, Sun Yat-sen University, Shenzhen, China; Shenzhen Key Laboratory for Systems Medicine in Inflammatory Diseases, Shenzhen Campus of Sun Yat-sen University, Sun Yat-sen University, Shenzhen, China; National Key Laboratory of Intelligent Tracking and Forecasting for Infectious Diseases, School of Medicine, Shenzhen Campus of Sun Yat-sen University, Sun Yat-sen University, Shenzhen, China

**Keywords:** Metatranscriptomics, Pathogen dicovery, Parasites, Small mammals, Infectome

## Abstract

Parasite communities in the internal organs of small mammals remain poorly characterized because metatranscriptomic pathogen discovery has largely focused on viruses and bacteria. We individually sequenced liver transcriptomes from 351 small mammals collected across 14 cities in Guangdong Province, China, and combined sequence-based detection with phylogenetic, abundance, transcriptional, and morphological analyses. Parasites were detected in 148 animals (42.2%). Four liver-associated taxa were identified, including three putatively novel or highly divergent lineages. Trichinellidae sp1 dominated the community, occurring in 36.2% of animals (127/351), five host species spanning Rodentia and Eulipotyphla, and all surveyed cities. *Hydatigera taeniaeformis* occurred in 3.7% (13/351), and its identification was supported by hepatic cysts and morphologically recovered strobilocerci. Eucoccidiorida sp1 and *Trypanosoma sp1* were detected sporadically. Mapping to available reference genomes revealed transcripts from multiple genes and functional categories, supporting genuine biological signals rather than isolated marker-gene matches. Amsterdam virus covaried with Trichinellidae sp1 (r = 0.59), whereas Wenzhou tapeworm virus 1 covaried with *H. taeniaeformis* (r = 0.47). These findings reveal an unexpectedly prevalent, phylogenetically diverse, and broadly distributed liver-associated parasite community and establish individual-organ metatranscriptomics as a useful framework for parasite discovery and surveillance.

**Author summary:** Rodents and shrews can host parasites of medical or veterinary importance, but their liver-associated parasites have rarely been surveyed systematically. We examined liver RNA from 351 small mammals collected throughout Guangdong Province. More than two-fifths carried at least one parasite. The most striking finding was an uncharacterized trichinellid parasite in over one-third of the animals. It occurred in five host species—including both rodents and shrews—and in every surveyed city, indicating an unusually broad host and geographic distribution. We also detected the tapeworm *Hydatigera taeniaeformis*, supported by visible liver cysts and recovered larvae, and two rare protozoan lineages. These findings reveal how much parasite diversity can remain hidden when surveillance overlooks particular organs or focuses on other pathogens. Individual-organ RNA sequencing can therefore complement conventional parasitology and help prioritize widespread or unusual lineages for taxonomic, ecological, and public-health investigation.

## Introduction

Small mammals, particularly rodents and eulipotyphlans, harbour a remarkable diversity of pathogens. Their close association with human settlements makes them important reservoirs of zoonotic agents (1, 2). Nevertheless, their parasites have received far less attention than viral and bacterial pathogens, despite the close links of many rodent-associated parasites to human disease. Representative examples include *Trichinella spp.*, which cause food-borne trichinellosis; *Hydatigera taeniaeformis*, whose larval stage develops in rodent livers; *Calodium hepaticum*, the causative agent of hepatic capillariasis; *Angiostrongylus cantonensis*, which causes eosinophilic meningitis; the occasionally zoonotic trypanosome *Trypanosoma lewisi*; and the tick-borne protozoan *Babesia microti*, for which rodents serve as important reservoirs (3–8). However, the diversity, prevalence, host range and geographical distribution of these parasites remain poorly characterized, particularly in subtropical regions where rodents frequently interact with humans, livestock and vectors.

Recent advances in metagenomic and metatranscriptomic sequencing have greatly accelerated pathogen discovery. However, these studies remain strongly virus-oriented, and the biological origins of sequences detected in vertebrate tissues are not always correctly resolved. Williams *et al.* initially identified two highly divergent negative-sense RNA viruses, Amsterdam virus and Fulton virus, in metagenomic data from wild house-mouse livers in New York City (9). Fluorescence *in situ* hybridization subsequently localized the viral RNA to adult *Calodium hepaticum* worms and their eggs, rather than to mouse hepatocytes, demonstrating that the viruses infected the parasite rather than the vertebrate host. Among 22 *C. hepaticum*-positive brown rats, Amsterdam virus, Fulton virus and both viruses were detected in 31.8%, 22.7% and 18.2%, respectively (9). These findings demonstrate that sequencing vertebrate tissues can reveal not only parasites but also microorganisms associated with the parasites themselves.

Indeed, metatranscriptomics can simultaneously profile viruses, bacteria, fungi and eukaryotic parasites, namely, the total “infectome”, as demonstrated by many studies of human respiratory disease, pigs, and companion cats and dogs (10–12). Its value for parasite discovery was particularly evident in a recent survey of small mammals in Guangdong Province, which analysed 2,408 lung, spleen and intestinal specimens from 858 rodents and shrews and identified 30 eukaryotic pathogen taxa, with an overall positivity rate of 40.8% (13). Importantly, the study uncovered multiple previously uncharacterized eukaryotic lineages, including putatively novel members of Giardia, Spironucleus, Brachylaima and Hepatozoon, highlighting the substantial unexplored parasite diversity in Guangdong small mammals. In a separate organ-specific investigation, Kuang *et al.* analysed kidney tissues from 142 bats and identified a previously undescribed apicomplexan protozoan, provisionally named *Klossiella yunnanensis*, in six of 20 pooled libraries (30%) (14). Its mitochondrial COX1 and cytB genes shared only 87.7% and 91.4% nucleotide identity with those of the closest recognized species, *K. equi*. These studies demonstrate the value of organ-based metatranscriptomics for discovering highly divergent parasites, even in the absence of closely related reference genomes.

Guangdong Province remains an important endemic area for human parasitic diseases, despite substantial declines in several major helminth infections over recent decades (15). Clonorchiasis, caused by the Chinese liver fluke *Clonorchis sinensis*, remains of particular concern. The parasite inhabits the hepatobiliary system of humans and other fish-eating mammals and is endemic across numerous counties in Guangdong (16). Provincial surveillance estimated a human prevalence of 4.25% in 2021, with the greatest disease burden concentrated around the Pearl River and Han River systems (17). Notably, the Guangdong survey by Xin *et al.* was restricted to lung, spleen and intestinal specimens and did not examine liver tissue (13), leaving the diversity of liver-associated parasites largely unexplored. Against this background, we used small-mammal liver specimens collected during clonorchiasis surveillance to perform individual metatranscriptomic sequencing of 351 animals from across Guangdong Province. We characterized the diversity, prevalence, host range, geographical distribution and transcriptional activity of liver-associated parasites, with particular attention to previously unrecognized taxa.

## Results

### Sample distribution and host identification

Between 2021 to 2025, we collected 351 small mammals from agriculture and residential setting of 14 prefecture-level cities across Guangdong Province (Fig. 1A). COX1-based host identification resolved nine named species and one putative novel lineage across five genera and two orders, Rodentia and Eulipotyphla. *Rattus norvegicus* the most frequently sampled species (n=174), followed by *R. tanezumi* (n=54), *Bandicota indica* (n=40), *R. andamanensis* (n=27), *Suncus murinus* (n = 23), and *R. losea* (n=20) (Fig 1B). Four animals formed a well-supported, divergent COX1 lineage within *Niviventer* that was distinct from *N. sacer* and *N. fulvescens* (Fig. 1B); we therefore provisionally designated this lineage *Niviventer sp*.

**Fig 1.**
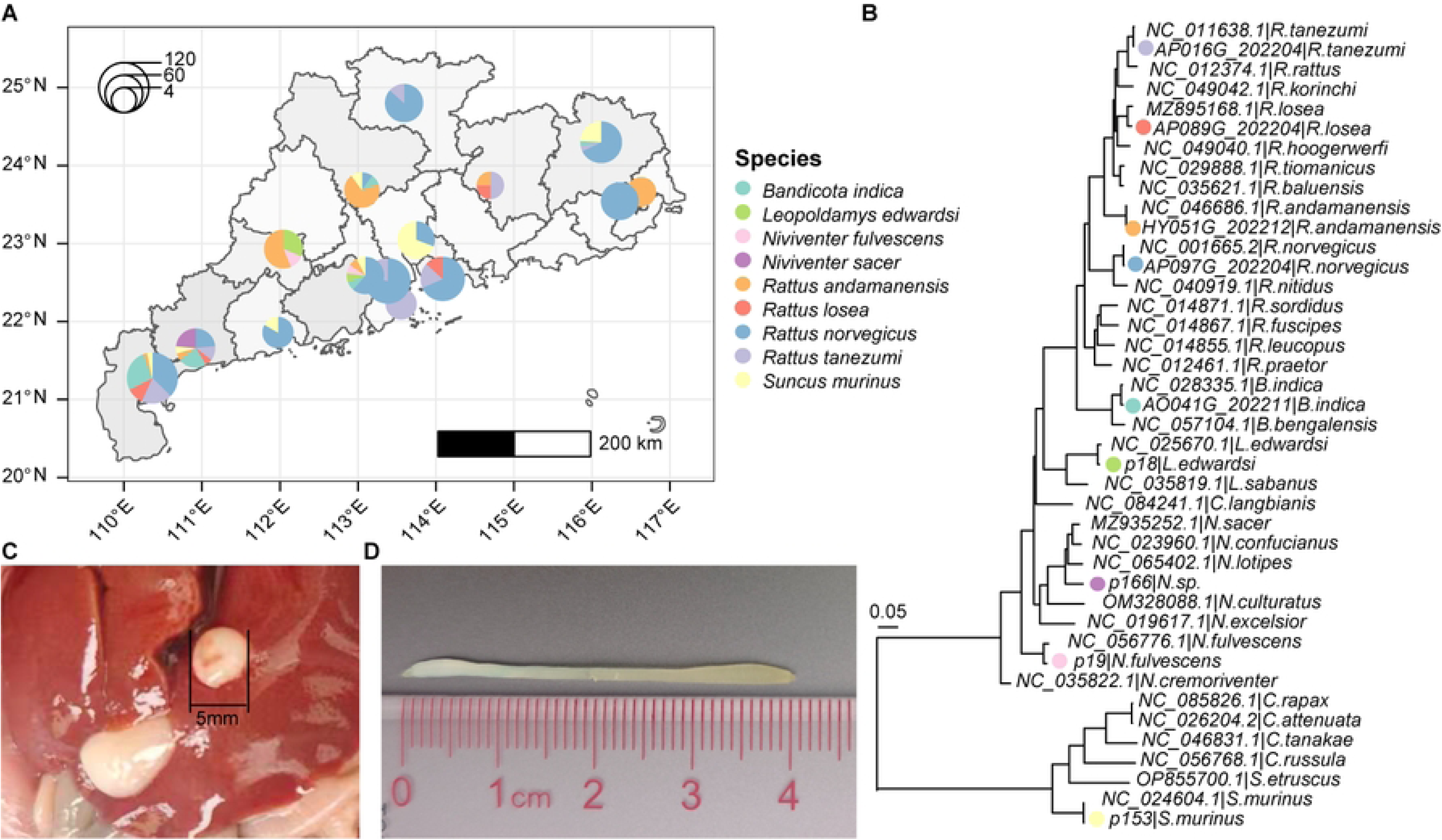
Sampling locations, host identification, and gross liver pathology. (A) Geographic distribution of the sampled small mammals in Guangdong Province. Pie charts show host-species composition at each location, and circle size is proportional to the number of animals sampled. Base map vector data were obtained from a publicly available dataset (https://geo.datav.aliyun.com/areas_v3/bound/440000_full.json) and visualized in ArcGIS. (B) Unrooted maximum-likelihood phylogeny inferred from full-length COX1 sequences. Colored circles indicate sequences generated in this study and correspond to the host colors in panel A. Branch lengths represent nucleotide substitutions per site. (C) Representative solitary or scattered milky-white cysts on the liver surface. (D) A strobilocercus recovered from a hepatic cyst.

Liver tissues were collected from all 351 animals and examined for gross lesions before sequencing (Table S1). Solitary or scattered milky-white cysts, ranging from several millimetres to several centimetres in diameter, were observed in some livers. Coiled, tape-shaped strobilocerci were recovered from these cysts (Fig 1C, D). These observations suggest the presence of parasites in these livers.

### Metatranscriptomic analyses reveal four liver-associated parasites

Individual metatranscriptomic sequencing of these liver tissue generated 351 libraries comprising 6.45 Tbp of data. Each library yielded a median of 93,135,802 reads (range, 937,552–579,603,958) (Table S1). Applying the detection criteria described in the Methods, we identified four liver-associated eukaryotic parasite taxa. Other than *Hydatigera taeniaeformis*, three represented previously uncharacterized species level taxa, including Trichinellidae sp1, Eucoccidiorida sp1, and *Trypanosoma sp1*.

To determine the taxonomic placement of the four parasites, we inferred a broad phylogeny from EF-1α protein sequences spanning Nematoda, Platyhelminthes, Apicomplexa, and Euglenozoa (Fig 2). The study sequences clustered within their expected higher-level groups: Trichinellidae, Taeniidae, Sarcocystidae, and Trypanosomatidae. These placements supported their family- or order-level assignments while revealing substantial divergence between several sequences reported here and their closest references.

**Fig 2.**
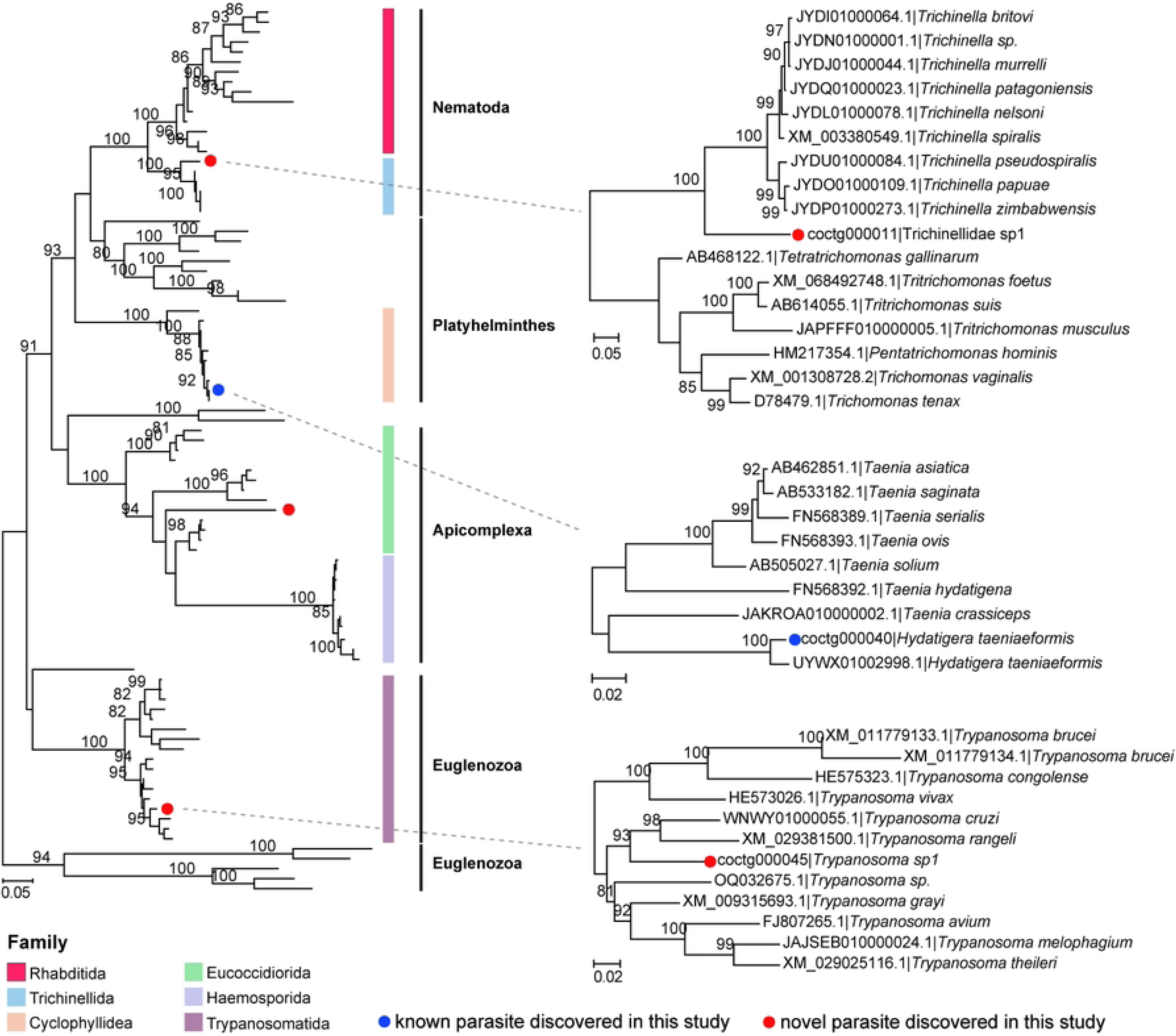
Detection and phylogenetic characterization of liver-associated eukaryotic parasites. The broad maximum-likelihood tree on the left was inferred from EF-1α amino acid sequences. The focused phylogenies of Trichinellidae, Taeniidae, and Trypanosoma on the right were inferred from EF-1α nucleotide sequences. Dashed lines connect these focused trees to their corresponding positions in the broad protein tree. Blue and red circles indicate known and putatively novel parasite species identified in this study, respectively. Trees were midpoint-rooted for visualization. Node labels indicate bootstrap support values, and scale bars represent substitutions per site.

*Hydatigera taeniaeformis* clustered with conspecific reference sequences and shared 100% EF-1α amino acid identity, supporting species-level identification. The remaining four parasites were provisionally designated as divergent species-level taxonomy. Trichinellidae sp1 formed a sister lineage to *T. zimbabwensis* and shared 94.2% amino acid identity with its closest reference. Eucoccidiorida sp1 was placed within the *Besnoitia* lineage but shared only 84.4% identity with available sequences. Trypanosoma sp1 shared 97.5% identity with *T. theileri* and fell within a clade containing *T. theileri*, *T. grayi*, and *T. avium*. Together, these analyses supported the provisional taxonomic assignments of the detected parasites and highlighted four lineages requiring further genomic characterization.

### Community structure, host range, and geographic distribution of liver parasites

Overall, 148 of 351 animals (42.2%) were positive for at least one parasite (Fig 3A, B). Of these, 142 animals (40.5% of all animals) carried one parasite and six (1.7%) carried two; no animal carried more than two parasite taxa. Each liver contained a mean of 0.4 parasite species (range, 0-2), with a mean total abundance of 2.75 RPM based on EF-1α protein gene (range, 0-71). The community was strongly dominated by Trichinellidae sp1, which occurred in 36.2% of animals (127/351), followed by *Hydatigera taeniaeformis* at 3.7% (13/351). Eucoccidiorida sp1 and *Trypanosoma sp1* were detected only sporadically (Fig. 3A).

**Fig 3.**
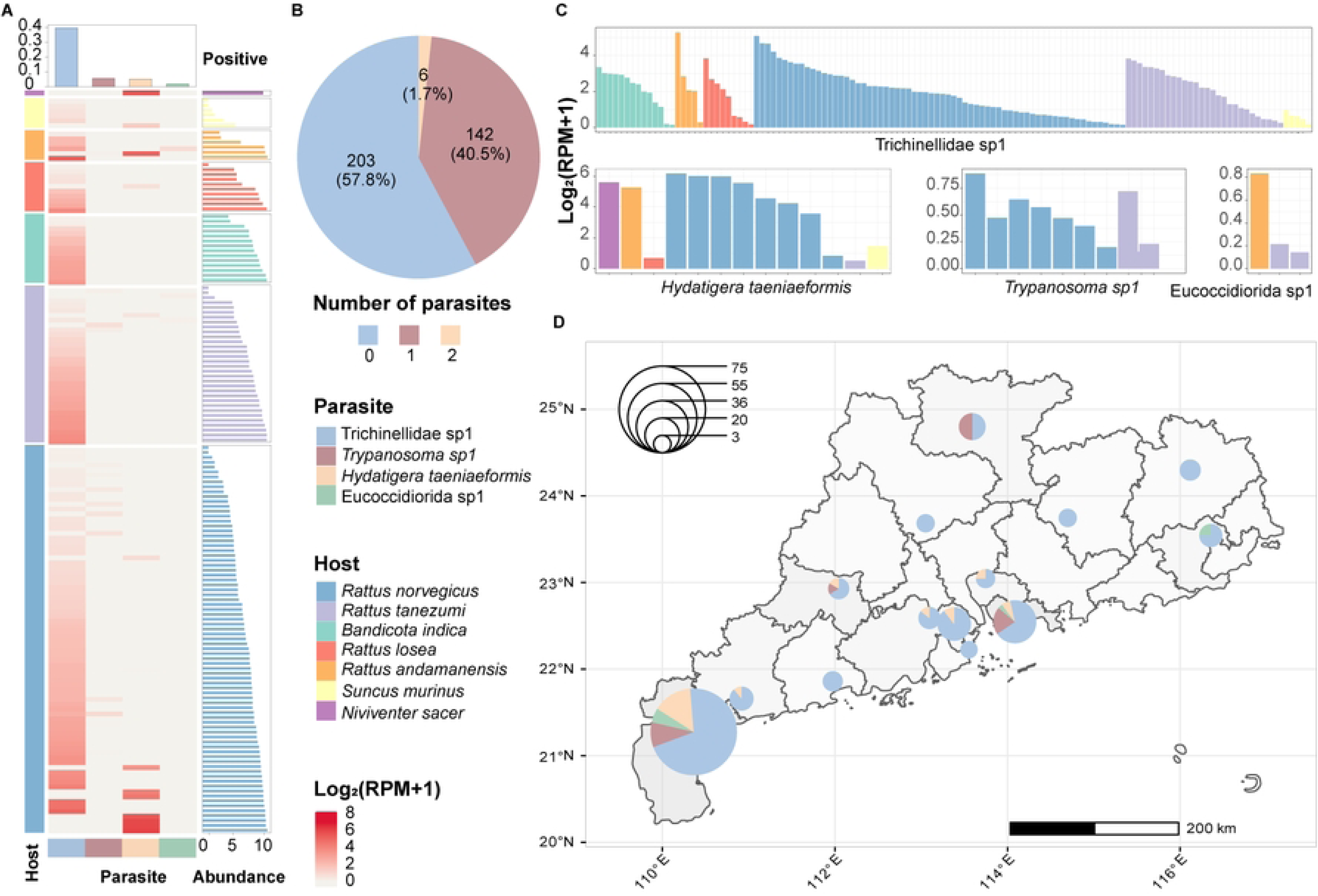
Community structure, co-infection patterns, host range, and geographic distribution of liver parasites. (A) Heatmap showing parasite abundance across individual animals. Rows represent animals and columns represent parasite taxa. The left annotation denotes host species, the top bars show parasite prevalence, and the right bars show total parasite abundance per animal. Heatmap values are displayed as log_2_(RPM + 1). (B) Proportions and numbers of animals carrying zero, one, or two parasite taxa. (C) Host-species composition of positive detections for each parasite. (D) Geographic distribution of the four parasites. Pie-chart colors indicate parasite composition at each location, and circle size is proportional to the number of parasite-positive animals.

Three of the four parasites occurred in at least two host species (Fig 3C). Trichinellidae sp1 had the broadest host range and was detected in *R. norvegicus*, *R. tanezumi*, *B. indica*, *Niviventer sp.*, and *S. murinus*. *Hydatigera taeniaeformis* occurred in four host species, whereas Eucoccidiorida sp1 and *Trypanosoma sp1* each occurred in two. Geographic distributions showed a similar contrast (Fig 3D). Trichinellidae sp1 was detected in all surveyed cities, whereas *H. taeniaeformis* was found in six cities, mainly in western Guangdong and the Pearl River Delta. *Trypanosoma sp1* was detected in four cities, and Eucoccidiorida sp1. Thus, the liver-parasite community was dominated by a widespread trichinellid lineage, while the remaining parasites exhibited more restricted host and geographic distributions.

### Transcriptional activity supports active liver infection by the detected parasites

To assess whether the detected parasite sequences reflected transcriptionally active infections, we mapped reads to available reference genomes or assembled transcriptomes and examined their gene-level coverage and functional profiles (Fig 4; S3-S4 Figs). For *H. taeniaeformis*, reads consistently covered hundreds of genes across positive samples, with broadly similar expression profiles among animals. The detected genes were associated with core cellular functions, including translation, cytoskeletal organization, energy metabolism, and protein processing (Fig. 4). Together with the morphological recovery of strobilocerci (Fig. 1C), these data support an active larval stage in the liver. Eucoccidiorida sp1 was detected at low abundance in only a few samples, but its reads mapped to multiple genes across the genome rather than to a single conserved region. The detected genes included functions related to translation, ribonucleoprotein complexes, nucleotide binding, and DNA replication, supporting genuine transcriptional activity (Fig. 4B).

**Fig 4.**
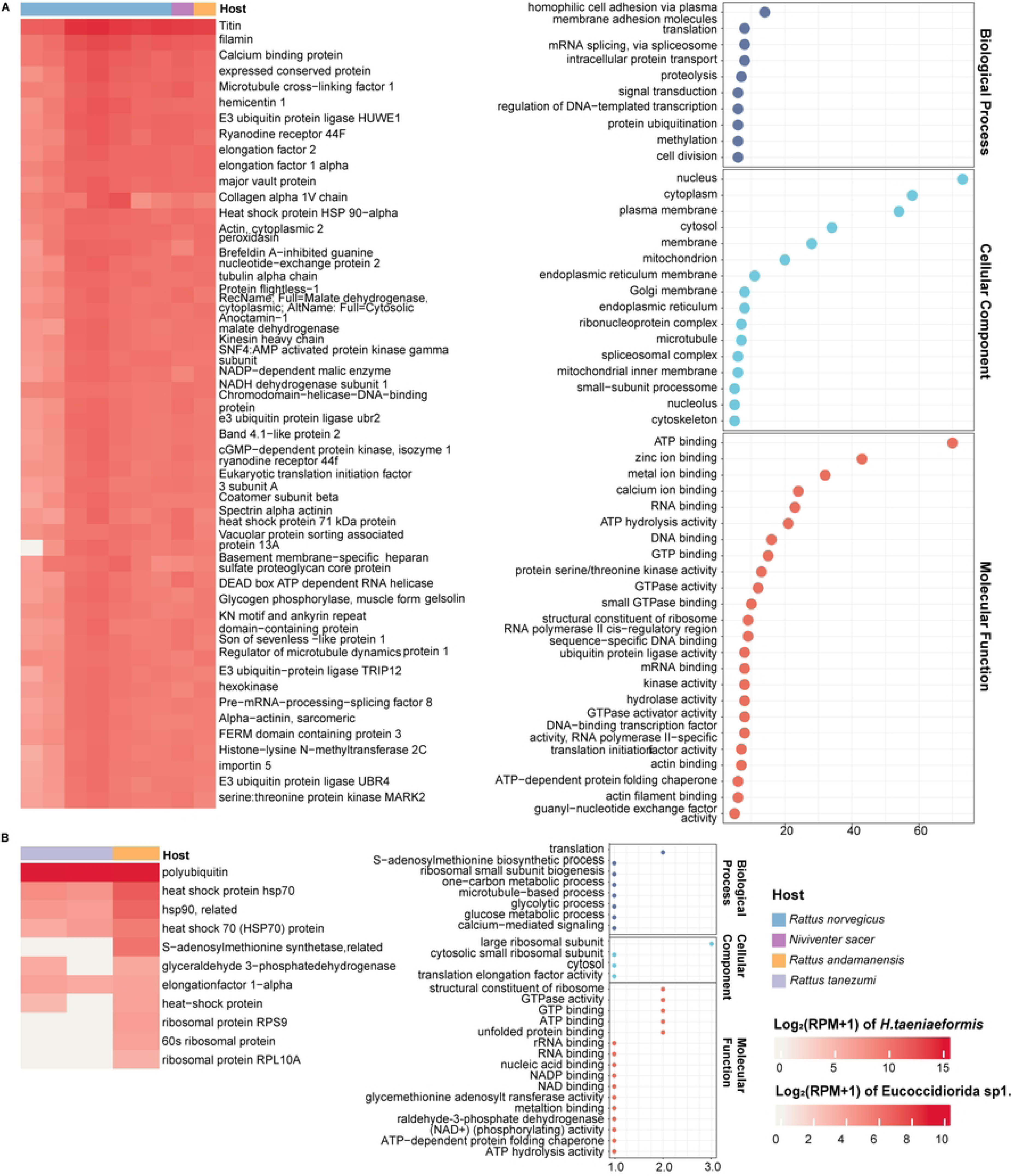
Gene-expression profiles and functional annotation of liver-associated parasites. The left panels summarize gene coverage and abundance across positive samples, with host species indicated by the top annotation bars. The right panels show Gene Ontology terms assigned to the detected genes, grouped into biological process, cellular component, and molecular function categories. (A) *Hydatigera taeniaeformis*. (B) Eucoccidiorida sp1.

### Parasites of the parasites: associations between invertebrate-associated viruses and liver parasites

In this study, we identified 25 viral species, most of which were phylogenetically related to viruses associated with invertebrate rather than vertebrate hosts. To investigate whether these viruses might be associated with liver parasites, we examined correlations between their abundances. This analysis identified two significant positive correlations, suggesting potential virus–parasite associations (Fig 5A, B). Amsterdam virus (*Usmuvirus newyorkense*) was positively associated with Trichinellidae sp1 (Pearson r = 0.59, p = 5.5e-16; Fig 5C), whereas Wenzhou tapeworm virus 1 (*Tapwovirus cesti*) was positively associated with *H. taeniaeformis* (r = 0.47, p = 0.026; Fig 5D). The remaining virus-parasite pairs showed weak associations. Guangdong *Bandicota indica* Jingchuvirales sp1 was detected in only one animal, precluding a robust association analysis (Fig 5A). RdRP phylogenies placed the study sequences of Wenzhou tapeworm virus 1 and Amsterdam virus within *Nyamiviridae* and *Lispiviridae*, respectively (Fig 5E; S5 Fig). Together, their abundance covariation and phylogenetic relationships support the hypothesis that these viruses are associated with the corresponding parasites rather than with the small-mammal hosts.

**Fig 5.**
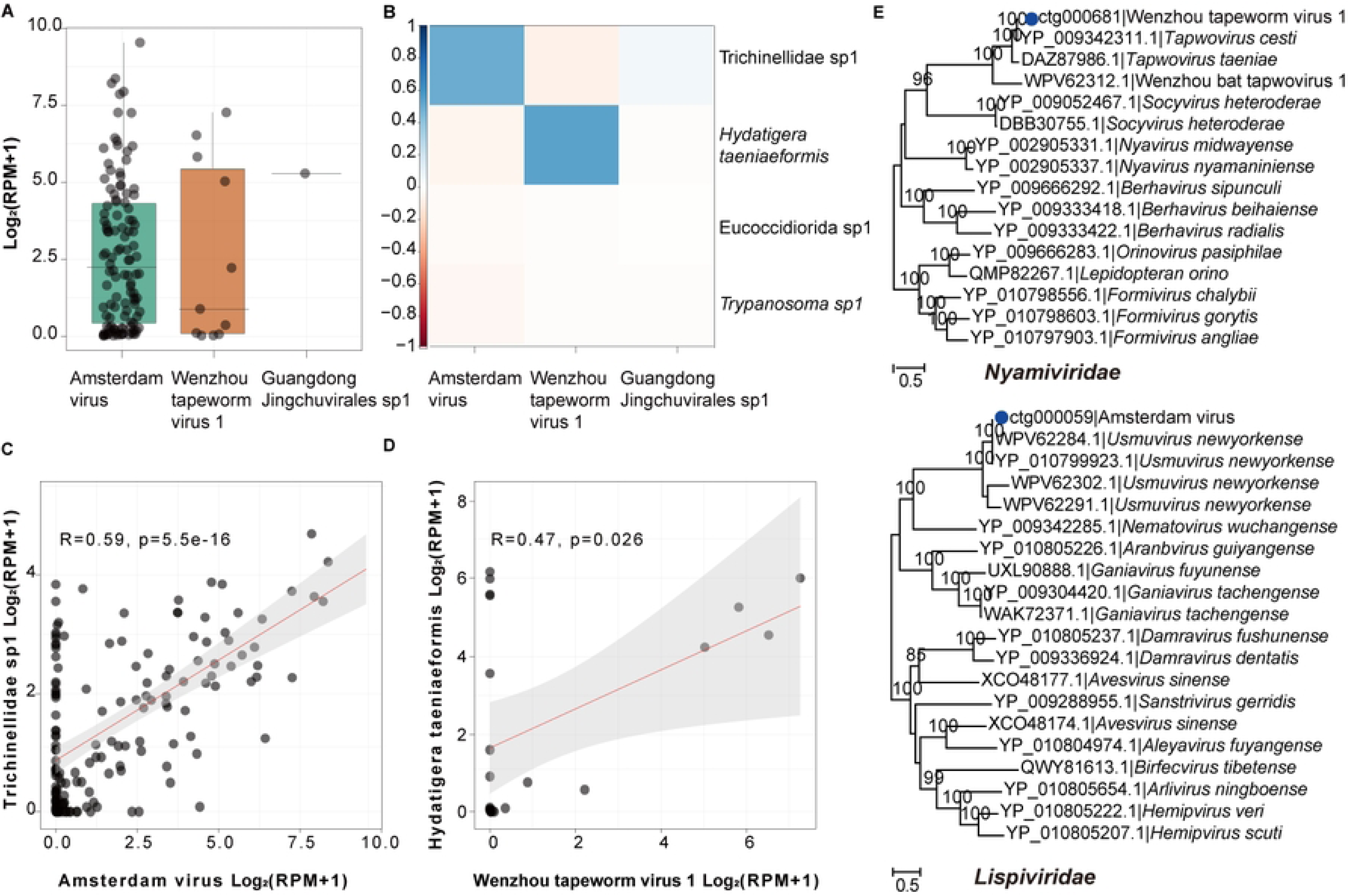
Associations between invertebrate-associated viruses and liver parasites. (A) log2(RPM + 1) abundances of the three invertebrate-associated viruses across positive libraries. (B) Pearson correlations between the three viruses and four parasites. Linear regressions between Amsterdam virus and Trichinellidae sp1 (C) and between Wenzhou tapeworm virus 1 and *Hydatigera taeniaeformis* (D). Shading indicates 95% confidence intervals; r and p values are shown. (E) Maximum-likelihood RdRp phylogenies of Wenzhou tapeworm virus 1 (upper) and Amsterdam virus (lower). Blue circles denote sequences identified in this study; node labels show bootstrap support and scale bars indicate amino acid substitutions per site.

## Discussion

In this study, individual-level metatranscriptomic sequencing revealed a diverse and largely uncharacterized assemblage of parasites in the livers of small mammals from Guangdong Province. Four parasite taxa were detected among 351 animals, four of which represented putatively novel or highly divergent lineages. Overall, 42.2% of the animals were positive for at least one parasite. The community was dominated by Trichinellidae sp1, detected in 36.2% of animals, whereas *Hydatigera taeniaeformis* occurred at a lower prevalence of 3.7% and the remaining protozoan lineages were sporadic. Together with the morphological recovery of strobilocerci and the detection of transcripts from multiple parasite genes, these findings demonstrate that small-mammal livers contain substantial parasite diversity that is not adequately captured by conventional surveillance.

The biological interpretation of these liver-associated parasites differs among taxa. The occurrence of *H. taeniaeformis* was expected because rodents are its principal intermediate hosts and its strobilocercus stage characteristically develops within hepatic cysts (18, 19). Indeed, a previous survey in northeastern China identified hepatic strobilocerci in 11.8% of brown rats (18). By contrast, the prevalence and abundance of Trichinellidae sp1 were unexpected. In the canonical *Trichinella* life cycle, adults reside in the intestine, newborn larvae disseminate through the circulation, and persistent larvae become established primarily in striated skeletal muscle (20). The liver is therefore not considered a major site of chronic *Trichinella* infection. Consequently, the Trichinellidae sp1 signal should not be interpreted simply as conventional hepatic trichinellosis. It may represent circulating larvae or parasite material within the hepatic vasculature, focal or ectopic tissue involvement, or a divergent trichinellid lineage with tissue tropism that differs from that of recognized *Trichinella* species. Its high prevalence and recurrent transcriptional signal nevertheless make incidental carry-over an insufficient explanation.

The detection of four parasite taxa in more than one host species also has important ecological implications. Trichinellidae sp1 had the broadest apparent host range, occurring in five small-mammal species spanning both Rodentia and Eulipotyphla. Multi-host parasitism is common among mammalian parasites, particularly when hosts share habitats, food resources, trophic interactions or arthropod vectors (21, 22). Closely related host species may also provide similar physiological environments, whereas trophic or vector-borne transmission can allow parasites to bridge greater host phylogenetic distances. The presence of the same parasite lineage in rodents and a shrew is therefore compatible with relatively weak host restriction, although the present cross-sectional data demonstrate a multi-host distribution rather than direct transmission events between these species. Comparative analyses have identified broad host range as a recurring characteristic of zoonotic and emerging pathogens (23, 24). Thus, the observed host breadth should be regarded as a surveillance-priority indicator: it increases the ecological opportunities for host encounters and adaptation.

The abundance correlations between several invertebrate-associated viruses and liver parasites provide an additional set of testable hypotheses. A relevant precedent is the discovery of Amsterdam and Fulton viruses in mouse-liver metagenomic data, followed by in situ localization of both viruses to adult *Calodium hepaticum* worms and their eggs rather than to mouse hepatocytes (25). Accordingly, the associations observed here are consistent with the possibility that some viruses infect the parasites themselves. However, correlation in bulk liver tissue cannot establish virus–parasite host assignment. Confirmation will require sequencing of individually isolated parasites, in situ hybridization or comparable spatial-localization approaches.

The absence of *Clonorchis sinensis* should also be interpreted in relation to its life cycle and the sampling design. Mammalian definitive hosts acquire *C. sinensis* by consuming freshwater fish containing metacercariae, after which the adult flukes migrate into and reside within the biliary system (26, 27). The sampled small mammals may have had limited access to infected fish, and their feeding ecology may therefore have provided few opportunities for exposure. In addition, sequencing a portion of liver parenchyma may miss parasites concentrated within focal intrahepatic bile ducts, particularly when infection intensity is low. The present result consequently indicates that *C. sinensis* RNA was not detected above the analytical threshold in the sampled liver tissues; it does not exclude infection in unsampled bile ducts or demonstrate that small mammals have no role in local transmission. Targeted examination of the bile ducts and gallbladder, fecal microscopy or PCR, and surveillance of fish-eating mammals would provide a more direct assessment of potential animal reservoirs.

The zoonotic implications of the detected parasites should therefore be considered cautiously but not dismissed. *H. taeniaeformis* is primarily maintained in a rodent–felid cycle, although exceptional human infections have been documented (28). The family Trichinellidae includes well-established food-borne zoonotic parasites, although the marked divergence and unusual tissue association of Trichinellidae sp1 mean that the zoonotic properties of named *Trichinella* species cannot automatically be assigned to this lineage (20). More generally, the widespread detection of several lineages across multiple small-mammal hosts suggests ecological flexibility and increases their priority for zoonotic assessment (23, 24). These findings do not demonstrate current transmission to humans. Instead, they identify a focused set of parasite lineages for proportionate follow-up based on phylogenetic proximity to human pathogens, host breadth, prevalence and plausible routes of human exposure.

Methodologically, our study demonstrates the utility of metatranscriptomics for parasite surveillance in animal organs. Metatranscriptomic sequencing has improved the detection and molecular characterization of blood parasites (29), while recent infectome studies have uncovered diverse known and novel eukaryotic pathogens in small mammals and highly divergent protozoa in bat kidneys (14, 30). We extend this approach to individually sequenced small-mammal livers and establish a practical workflow for future investigations. Briefly, reads remaining after host subtraction were assembled separately for each animal; candidate parasite contigs were identified through nucleotide- and protein-level similarity searches, evaluated by read remapping and sequence coverage, and placed phylogenetically using conserved marker genes, with transcripts from multiple parasite genes providing complementary support for their presence. Whenever suitable material is available, sequence-based evidence should be corroborated by morphological examination, targeted PCR, or histopathology. Such an evidence-integrated approach is particularly important for highly divergent eukaryotes, whose taxonomic assignments may otherwise be imprecise or misleading because of the limited and uneven representation of parasites in reference databases.

Several limitations should be acknowledged. First, only liver tissue was examined, preventing direct comparison with blood, skeletal muscle, intestine, bile ducts and other organs that may contain different parasite stages. Second, RNA-based detection depends on transcriptional activity and RNA preservation; absence of detectable RNA therefore does not establish absence of infection. Third, most novel lineages were classified primarily using EF-1α and transcriptome-derived sequences, which are sufficient for provisional phylogenetic placement but not formal species delimitation. Fourth, histological localization and targeted molecular confirmation were not available for every taxon, and RPM values cannot be interpreted as absolute parasite burdens (31). Finally, the low number of detections for several protozoa, uneven host and geographic sampling, and the cross-sectional design limit inference regarding prevalence, transmission direction, host competence and zoonotic potential.

In summary, the small-mammal liver represents an informative but underexplored compartment for parasite discovery. The unexpectedly widespread Trichinellidae sp1 and the morphologically and molecularly confirmed *H. taeniaeformis* collectively reveal a parasite community with substantial biological and potential public-health relevance. Although the present findings do not establish zoonotic transmission, their phylogenetic relationships and broad host distributions provide a clear rationale for targeted One Health surveillance.

## Methods

### Ethics statement

All procedures involving wild small mammals were reviewed and approved by the Ethics Committee of Sun Yat-sen University (SYSU-IACUC-MED-2021-B0123) and the Ethics Review Subcommittee for Animal Experiments and Human Trials of Health-Related Products, Guangdong Provincial Center for Disease Control and Prevention (20250032).

### Sample collection

Between 2021 and 2025, 351 small mammals representing the orders Rodentia and Eulipotyphla were collected from 14 prefecture-level cities in Guangdong Province, China. Sampling locations, collection dates, and host metadata are provided in S1 Table. Sampling at each locality was conducted over 3–4 consecutive days, with the duration adjusted according to weather conditions, particularly rainfall. On each trapping day, 200 baited live traps were deployed in agricultural areas and another 200 in residential areas. Traps were baited with a mixture of sweet potato, fried dough sticks, and cooked meat. At residential sites, 2–3 traps were placed near the corners of each selected farmhouse. At agricultural sites, 2–5 traps were positioned in brushwood or gullies at each trapping point, with the number of trapping points adjusted according to field size. Traps were set at dusk and checked at sunrise the following morning. Captured animals were euthanized and dissected, and multiple tissues were subsequently collected. Before tissue sampling, each liver was examined macroscopically, and visible lesions were documented and photographed. Representative hepatic cysts were opened, and the recovered strobilocerci were examined and photographed. The collected liver tissues were immediately immersed in RNA Stabilization Solution (Thermo Fisher Scientific, USA), transported on dry ice, and stored at −80°C until processing.

### RNA extraction, library preparation, and sequencing

Each of the 351 liver samples was processed individually. Approximately 50 mg of liver tissue was homogenized in 600 μL of lysis buffer using a TissueRuptor homogenizer (Qiagen, Germany). Total RNA was extracted using the RNeasy Plus Mini Kit (Qiagen, Germany) according to the manufacturer’s instructions. Sequencing libraries were prepared using the VAHTS Universal V8 RNA-seq Library Prep Kit for Illumina (Vazyme, China, Cat# NR605). The generated libraries were sequenced on the DNBSEQ-T series platform (MGI Tech, China) to generate 150-bp paired-end reads, with a target yield of 10 Gbp per sample.

### Processing of sequencing data

Adapter sequences and low-quality reads were removed from each library using fastp v0.20.1 (parameters: -q 20 -n 2 -y -c -p -G) (32). Residual rRNA reads were identified by mapping against the SILVA rRNA database (release 138.1) (33, 34) with Bowtie2 (version 2.5.4) (33, 34) and were discarded. Duplicate and low-complexity reads were removed using bbduk tool from the BBMap suite (35).

### Molecular identification of small-mammal hosts

Host species were identified from mitochondrial COX1 sequences recovered from the metatranscriptomic assemblies. Open reading frames were predicted with NCBI ORFfinder, and candidate COX1-containing contigs were identified by BLASTn v2.14.1 searches against COX1 reference sequences in NCBI RefSeq (E-value ≤ 1 × 10⁻¹⁰) (36, 37). The cytochrome c oxidase subunit I conserved domain (cd01660) was confirmed by RPS-BLAST searches against the NCBI Conserved Domain Database (38). Reads were mapped back to each candidate COX1 contig with Bowtie2 to inspect coverage and exclude poorly supported or chimeric assemblies. Study sequences were analysed together with representative reference sequences in a maximum-likelihood phylogeny inferred with IQ-TREE (version 1.6.12) (39). Final host assignments were based on both sequence similarity and phylogenetic placement.

### Discovery and taxonomic assignment of eukaryotic pathogens

A curated reference database of eukaryotic elongation factor 1-alpha (EF-1α) protein sequences was constructed from parasite sequences downloaded from NCBI. Redundant sequences were removed with CD-HIT (40), and the remaining sequences were subjected to preliminary phylogenetic analysis. Reference sequences with inconsistent annotations or anomalous phylogenetic positions were removed after manual review. Clean reads from each library were searched against this database using DIAMOND BLASTx v2.0.14 (E-value ≤ 1 × 10⁻⁵) (41). Matching reads were extracted separately for each sample and assembled with metaSPAdes (v4.2.0) (42). The resulting contigs were searched against the NCBI nucleotide and non-redundant protein databases using BLASTn (E-value ≤ 1 × 10⁻¹⁰) and BLASTx (E-value ≤ 1 × 10⁻⁵), respectively. Candidate parasite contigs were retained only when similarity searches and phylogenetic placement yielded concordant taxonomic assignments and the quantitative detection criteria described below were met. Sequences that formed distinct, well-supported lineages and could not be assigned confidently to a described species were given provisional labels based on the nearest supported higher-level taxon (for example, Trichinellidae sp1); these labels do not constitute formal taxonomic descriptions.

### Discovery of viruses

Clean non-rRNA reads were assembled into contigs using MEGAHIT (version 1.2.8) with default parameters and a minimum contig length of 300 bp(43). These contigs were searched against the NCBI non-redundant protein database using Diamond BLASTx (E-value ≤ 1 × 10⁻⁵). Contigs with viral matches were extracted and re-examined against the NCBI nucleotide and RefSeq genome databases using BLASTn (E-value ≤ 1 × 10⁻¹⁰) to remove host-derived or other non-viral regions. Viral identity was further assessed by the presence of supergroup or family-appropriate marker proteins, including RdRP for RNA viruses, NS1 for Parvoviridae, hexon for Adenoviridae, capsid protein for Circoviridae, and ORF1 for Anelloviridae. Putative host association was inferred from phylogenetic placement and the host metadata of the closest known relatives and was regarded as provisional in the absence of experimental validation.

### Quantification and detection criteria

To quantify taxon abundance, clean reads were mapped in end-to-end mode with Bowtie2 to representative viral genomes or to validated parasite marker sequences. Abundance was expressed as reads per million non-rRNA reads (RPM), calculated as the number of mapped reads divided by the total number of non-rRNA reads in the library and multiplied by 10⁶. To reduce false-positive detections caused by index hopping, a taxon was excluded from a library when its read count was less than 0.1% of the highest read count observed for that taxon in any library from the same sequencing lane. A taxon was considered positive only when mapped reads covered at least 300 nt of its marker gene and its abundance was ≥ 1 RPM. Prevalence was calculated as the proportion of the 351 animals meeting these criteria. Parasite richness was defined as the number of parasite taxa detected in a sample.

### Phylogenetic analyses

Representative sequences related to the study taxa were retrieved from NCBI/GenBank. For parasites, a broad phylogeny was inferred from EF-1α amino acid sequences spanning Nematoda, Platyhelminthes, Apicomplexa, and Euglenozoa, and focused nucleotide phylogenies were inferred for Taeniidae, and *Trypanosoma*. Viral phylogenies were inferred from the relevant marker proteins, including RNA-dependent RNA polymerase. Sequences were aligned with the L-INS-i algorithm in MAFFT v7 (44), and ambiguously aligned regions were removed with trimAl (45). Maximum-likelihood trees were inferred with IQ-TREE v3.0, with the best-fitting substitution model selected by ModelFinder. Branch support was evaluated using 1000 bootstrap. Trees were midpoint-rooted for visualization. Pairwise amino acid or nucleotide identities were calculated from the corresponding trimmed alignments.

### Assessment of parasite transcriptional activity

For parasite taxa with suitable reference genomes, including *Hydatigera taeniaeformis* and Trichinellidae *sp1*, clean reads from positive libraries were mapped to the closest available annotated genome using Bowtie2. Gene-level coverage and abundance were calculated from the resulting alignments. We performed functional annotation via sequence homology alignment as described above, and then extracted the corresponding GO terms for each gene using Blast2GO (v6.0). GO enrichment analysis was conducted using the clusterProfiler package (v4.4.4) in R, employing the over-representation analysis (ORA) method. Based on the annotation results, a customized gene–GO annotation file was used for the analysis. The background gene set (Universe) was defined as all genes that received at least one GO annotation. Significantly enriched GO terms were identified using a Benjamini– Hochberg-adjusted P-value (p.adjust) < 0.05 as the significance threshold. Transcriptional activity was considered broadly supported when reads mapped to multiple non-overlapping genes distributed across the reference genome and represented several functional categories, rather than being confined to EF-1α or another single conserved marker.

### Parasite community, host range, and geographic analyses

Co-infection was summarized as the number of parasite taxa detected in each animal. Host range was defined as the number of molecularly identified host species in which a parasite met the positivity criteria, and geographic range was summarized at the city level. Maps were produced in ArcGIS using the Guangdong administrative-boundary dataset available at https://geo.datav.aliyun.com/areas_v3/bound/440000_full.json.

### Virus-parasite association analysis

Potential associations between three invertebrate-associated viruses and the four liver parasites were evaluated using log2(RPM + 1)-transformed abundances. For each virus-parasite pair, the analysis included only libraries positive for at least one member of the pair were included. Pearson correlation coefficients and two-sided nominal P values were calculated, and selected associations were visualized by linear regression with 95% confidence intervals. These analyses were exploratory and were interpreted together with viral phylogenetic evidence; correlation alone was not considered proof of a virus-parasite host relationship.

## Data availability

Raw metatranscriptomic reads are available in the China National Genebank (CNGB) under BioProject ID PRJCA072689. The sample metadata, abundance matrices, alignments, phylogenetic trees, and analysis code required to reproduce the reported findings are available in Figshare under the link https://doi.org/10.6084/m9.figshare.33390832.

## Author contributions

Conceptualization, *D.W.*, *J.-D.C.*, and *M.S.*; Methodology, *M.-W.P.* and *M.S.*; Investigation, *M.-W.P.*, *D.W.*, and *M.S.*; Sampling and sample processing, *M.-W.P., Q.-Q.C., Y.-Q.L., Q.M., L.-J.C., J.-H.L.*; Writing – Original Draft, *M.-W.P.,* and *M.S.*; Writing-Review and Editing, All authors; Funding Acquisition, *J.-D.C, D.W.*, and *M.S.*; Resources (sampling), *M.-W.P., Q.-Q.C., G.-Y.X, D.W.* and *W.-C.L*; Resources (Computational), *J.-D.C, D.W*, and *M.S*.; Supervision, *D.W.*, *J.-D.C.* and *M.S.*.

## Acknowledgement

This work was supported by Prevention and Control of Emerging and Major Infectious Diseases-National Science and Technology Major Project (2025ZD01901103 and 2025ZD01901104). Talent Support Program of Guangdong Provincial Center for Disease Control and Prevention (2024D351). We gratefully acknowledge the staff of the Guangdong Provincial Center for Disease Control and Prevention for their valuable assistance with sample collection and members of our research team for their support with laboratory work.

## Notes

### Competing Interest Statement

The authors have declared no competing interest.

